# Spatiotemporal expression of *FRIGIDA* modulate flowering time in *Arabidopsis thaliana*

**DOI:** 10.1101/467613

**Authors:** Xiangxiang Kong, Jinjie Zhao, Landi Luo, Qian Chen, Guanxiao Chang, Jinling Huang, Yongping Yang, Xiangyang Hu

## Abstract

*FRIGIDA (FRI)* as the major regulator of flowering time in *Arabidopsis* accessions can activate its target *FLOWERING LOCUS C (FLC)* to delay flowering before vernalization. Besides *FLC*, other *FRI* targets also exist in *Arabidopsis*. Although leaves sense environmental cues to modulate flowering time, it is not known if roots also regulate the floral transition. In this study, we investigated the spatiotemporal effect of *FRI* on flowering time. Local expression of *FRI* in the phloem and leaves activated *FLC* to delay flowering. Furthermore, we found that local expression of *FRI* in the roots also delayed flowering by activating other targets *MADS AFFECTING FLOWERING4 (MAF4)* and *MAF5* in the roots. Graft and genetic experiments revealed that the spatial expression of *FRI* in the root might generate a mobile signal, which is transmitted from roots to shoot and antagonizes the *FT* signal to delay flowering. Specifically expressing *FRI* in the embryo efficiently delayed flowering, even expressing *FRI* as early as pro-embryo stage is enough to upregulate *FLC* expression to delay flowering. Together, our findings confirm the spatiotemporal effect of *FRI* on delaying flowering, and propose that root tissue also perceives the flowering signal to fine-tune the flowering time through *MAF4/5* as novel targets of *FRI*.

**Highlight:** Root FRIDIGA activated the novel targets *MAF4/5* to delay flowering; Temporal expressing *FRIGIDA* at as early as pro-embryo stage is efficient to delay flowering.

## Introduction

Flowering at the appropriate time is fundamental to reproductive success of plants and their adaptation to environmental stress. As such, plants have evolved accurate mechanisms to initiate flowering in response to exogenous environmental cues or endogenous signals (Jaeger *et al*., 2006; Kim *et al*., 2009; Amasino and Michaels, 2010). Many plant species require a long period of cold treatment (i.e., vernalization) before flowering (Kim *et al*., 2009). *FRIGIDA (FRI)* is a major regulator of flowering time in various *Arabidopsis thaliana* accessions (Johanson *et al*., 2000). The winter-annual (late flowering) and rapid-cycling (early flowering) accessions are often distinguished by allelic variation at *FRI* and *Flowering Locus C (FLC)* (Shindo *et al*., 2005; Shindo *et al*., 2006). *FRI* encodes a novel protein with two potential coiled-coil domains that activates the expression of *FLC*, which in turn encodes a MADS box transcriptional factor that quantitatively inhibits the floral transition through repressing the flowering pathway integrators *FLOWERING LOCUS T* (*FT*) and *SUPPRESSOR OF OVEREXPRESSION OF CONSTANS1* (*SOC1*) in *Arabidopsis* (Johanson *et al*., 2000; Michaels, 2004; Schmitz, 2007).

Although *FRI* was isolated two-decade ago, its biochemical function and the mechanism underlying flowering regulation are largely unknown. FRI acts as a scaffold protein that interacts with FRL1, SUF4, FLX, FES1, UBC1 and CBP20 to form a transcription activator complex, which recruits chromatin modification factors, such as the *SWR1* complex and *SET2* homolog, to epigenetically modify the histone methylation level at the *FLC* locus (Choi *et al*., 2011; Li *et al*., 2018). Besides *FLC*, other five *FLC* homologues, including *FLOWERING LOCUS M (FLM)/MADS AFFECTING FLOWERING 1*, and *MAF2*-*MAF5*, also modulate flowering time (Ratcliffe *et al*., 2003; Scortecci *et al*., 2003; Gu *et al*., 2009; Kim *et al*., 2010). *MAF4* can directly interact with other *FLC* homologues proteins to repress *FT* expression (Gu *et al*., 2013). *MAF4* and *MAF5* is rather specific to preventing a precocious vernalization response (Kim and Sung, 2013). Nevertheless, the underlying mechanisms remain to be discovered.

In many plants, the transition from vegetative to reproductive development is controlled by environmental cues, such as day length and temperature (Amasino and Michaels, 2010). The environmental signal is usually perceived by the leaves, while flowers develop from primordia formed on the flanks of the SAM (shoot apical meristem). Ectopic expression of *FLC* in the phloem of leaves can reduce *SOC1* and *FT* mRNA levels to delay flowering (Searle *et al*., 2006). Unlike these findings in leaves, previous studies show that ectopic expression of *FLC* in the root did not efficiently delay the flowering time (Searle *et al*., 2006), and ectopic expression of *FT* in the root also could not efficiently promote flowering time (Abe *et al*., 2005), casting doubt on the contribution of the root to modulating flowering time. However, other lines of evidence imply that the root tissue might regulate flowering time. For example, in white mustard (*Sinapis alba*), a shoot-to-root-to-shoot physiological loop driving sugar and cytokinin fluxes is essential for flowering (Bernier *et al*., 1993). Data mining of public microarray experiments showed that about 200 genes associated with flowering time are expressed in the root (Bouche *et al*., 2016). Thus, studying the spatial and temporal effects of *FRI* on flowering time should reveal at which developmental stage *FRI* epigenetically regulates gene expression to delay flowering and whether root tissue modulates flowering time.

In this study, we systemically investigated the effect of various spatial and temporal expression patterns of *FRI* on flowering time in *Arabidopsis*. We found that expression of *FRI* in the leaves or phloem activated *FLC* transcription in a cell-autonomous manner and thereby delayed flowering. In contrast to *FLC*, which does not delay flowering when spatially expressed in root tissue (Searle *et al*., 2006), we found that ectopic expression of *FRI* in the roots also delayed flowering by activating the transcription of *MAF4* and *MAF5*, and subsequently suppressed the downstream *FT* signal depending on downstream mobile signals, such as antiflorigen to delay flowering. Furthermore, expressing *FRI* only during embryonic development or in the young seedling stage using temporal-specific promoters also efficiently delayed flowering time, and was accompanied by high levels of histone methylation at the *FLC* locus. Based on our data, we propose that, in addition to leaves, which are believed to be the main sensors of the environmental signal, roots also play critical roles in modulating flowering time. Thus, our data provide novel insight into the molecular mechanism underlying *FRI*-mediated regulation of flowering time.

## Materials and methods

### Plant materials and growth conditions

Wild-type *Arabidopsis thaliana* (Col) plants were used in this study. Col seeds were sown in plastic pots under a 16-h light/8-h dark photoperiod. For vernalization, seeds were germinated and pre-grown for 7 days under standard conditions (16 h light/ 8 h darkness, 22°C). The plants were transferred to cold conditions (8 h light /16 h darkness, 4°C) for 4 weeks, and then returned to standard conditions. The time to flowering was determined as the total number of rosette leaves when the floral bolt was 1 cm high (Sheldon *et al*., 2008; Hu *et al*., 2014).

### Development of transgenic lines

To construct the *35S:GFP*-*FRI* plasmid, the full-length cDNA coding region of *FRI* was amplified and cloned downstream of the *GFP* fluorescence marker in the *pEGAD* vector. To construct the *35S:GUS*-*FRI* plasmid, the coding region of *GFP* of *35S:GFP*-*FRI* was replaced by the *GUS* fragment. To generate the *FRI::GUS*-*FRI* construct, the promoter *FRI* with 2.1 kb of genomic sequence were amplified from winter-annual accession *H51*, and the 35S promoter of the *35S::FRI*-*GUS* construct was replaced with the *FRI* promoter. To drive *FRI* expression by the spatiotemporal-specific promoters, the promoters *SUC2*, *ML1*, *KNAT*, *RolC*, *TobRB7*, *LEC2*, and *EM1* were amplified, and the 35S promoter of the *35S::FRI*-*GUS* construct was replaced with the above promoters, respectively. To construct the *TobRB7::MAF4* or *TobRB7::MAF5* plasmid, the full-length cDNA of *MAF4* or *MAF5* was amplified and replaced the *GFP* fluorescence marker in the *pEGAD* vector, then the *35S* promoter was replaced with the *TobRB7* promoter. To generate the inducible *ProER8:GUS*-*FRI* plasmid, the coding region of *GUS*-*FRI* was amplified and inserted into the *pER8* vector. These constructs were directly transformed into *Agrobacterium tumefaciens* GV3101, and the modified floral dip method was performed to generate the corresponding transgenic plants in the Col background. The transgenic seeds were screened on half-strength Murashige and Skoog medium containing 10 mg/L Basta or 35 mg L^-1^ Hygromycin. The T3 homologous transgenic seeds were used in our experiments. All of the primer sequences are listed in Supplemental Table 1.

### RNA extraction and quantitative RT-PCR

Total RNA was extracted using Trizol reagent (Invitrogen). Quantitative real-time PCR (qRT-PCR) was performed as described previously. Briefly, first-strand cDNA was synthesized from 1.5 μg DNAse-treated RNA in a 20-μL reaction volume using M-MuLV Reverse Transcriptase (Invitrogen) with oligo (dT)_18_ primer. RT-PCR was performed using 2XSYBR Green I Master on a Roche LightCycler 480 real-time PCR machine, according to the manufacturer’s instructions. At least three biological replicates for each sample were used for the RT-PCR analysis, and at least three technical replicates were analyzed for each biological replicate. The *Actin* gene was used as a control. Gene-specific primers used to detect transcripts are listed in Supplemental Table 1.

### ChIP assay

ChIP was performed largely as described previously (Hu *et al*., 2014). Immunoprecipitation was performed with acetylated H3 (1:1000; Millipore) and trimethyl H3K4 (1:500; Millipore) or trimethyl H3K27 (1:500; Millipore) antibody Both immunoprecipitated DNA and input DNA were analyzed by real-time PCR. Primers and PCR detection of *FLC* regions were as described (in Supplemental Table 1). Data from ChIP experiments are expressed as means ±SD of three biological replicates.

### Protoplast transient assay

Rosette leaves of *Arabidopsis* Col plants that had been grown for 4 weeks under long-day conditions (16 h light/8 h darkness) were sampled for the isolation and transformation of protoplasts as described (Yoo *et al*., 2007). All of the plasmid DNA for the protoplast transformation was prepared by the CsCl gradient method. Twelve hours after transformation, the protoplasts were observed. Fluorescence of the GFP chimeric gene was detected using an Olympus FluoView confocal microscope with excitation and emission filters of 450–490 nm and 520–560 nm, respectively.

### Histochemical analysis of GUS activity

Whole tissues were vacuum-infiltrated in GUS-staining buffer [0.5 mM NaPO_4_ (pH 7.0), 10 mM EDTA, 1 mM potassium ferrocyanide, 1 mM potassium ferricyanide, 1 mM 5-bromo-4-chloro-3-indolyl-ß-d-glucuronic acid (X-Gluc), and 0.1% Triton X-100], incubated at 37°C, and then destained in 70% ethanol. Cross-sections of leaf and stem tissues were prepared using a razor blade followed by staining as described above. For cross-sections of cotyledons, GUS-stained seedlings were fixed (50% ethanol, 5% acetic acid, 3% formaldehyde), dehydrated in a graded ethanol series (70%, 96%, and 100%; 2 h each), embedded in Technovit 7100 (Heraeus Kulzer), according to the manufacturer’ s instructions, and sectioned using a microtome.

### Immunoblot assay

Total proteins were prepared by grinding seedlings on ice in extraction buffer (50 mM Tris, 5% glycerol, 4% SDS, 1% polyvinylpolypyrrolidone, 1 mM phenylmethylsulfonylfluoride (pH 8.0)), followed by centrifugation at 4°C and 14,000 g for 15 min. A 15-μg aliquot of protein was separated by electrophoresis on a 12% SDS–polyacrylamide gel and blotted onto polyvinylidene difluoride membranes, which were then probed with the appropriate primary anti-GFP (1:3000, Clontech) or anti-actin (1:1000, Sigma-Aldrich) antibody and horseradish peroxidase-conjugated goat anti-mouse secondary antibody (1:3000, Promega). Signals were detected using the ONE-HOUR IP-Western Kits (Cat. L00232, Genescript).

### Grafting methods

The general grafting conditions and procedures were performed as reported previously (Turnbull *et al*., 2002). Briefly, young seedlings with the cotyledons expanded were used as the scions and rootstocks. We used a sapphire/diamond knife to remove the cotyledons from all seedlings, and cut the scion and rootstock seedlings cleanly across the hypocotyl just below the cotyledon stumps. The rootstock and scion were well-matched for size where the cut ends could be pressed very closely together. After all grafts were complete, the grafted plants were placed in petri dishes in the growth room for 3-5 days, and then were moved to soil and grown until flowering, and the flowering time was scored.

### Accession numbers

The T-DNA insertion lines were used in this study: *maf4*-*1(salk*_*028506)*, *maf4*-*1(salk* _*095092)*, *ft*-*10* (*CS9869*) and *flc*-*3* (*Germplasm:1008704442*)

## Results

### Constitutive expression of *FRI* delays flowering time

*FRI* upregulates *FLC* transcription and thereby delays flowering time (Geraldo *et al*., 2009; Choi *et al*., 2011). To decipher the role of *FRI* in controlling flowering time, we investigated the expression pattern of *FRI* using constructs in which the *GUS* reporter marker is driven by the endogenous *FRI* promoter. As shown in Fig. 1A, *GUS* staining could be observed in the young embryo and cotyledon, and in the leaves, vasculature, meristem, and roots of 2-week-old seedling. We then generated a construct containing *FRI* fused to green fluorescence protein (*GFP*) under the control of a cauliflower mosaic virus *(CaMV) 35*S promoter (termed *35S::GFP*-*FRI*). When *35S::GFP*-*FRI* was transiently expressed in *Nicotiana benthamiana* leaves or in protoplasts derived from *Arabidopsis* Col leaves, fluorescence was detected only in the nucleus (Fig. 1B), which is consistent with our previous results that *FRI* is mainly localized in the nucleus (Hu *et al*., 2014).

**Fig. 1.**
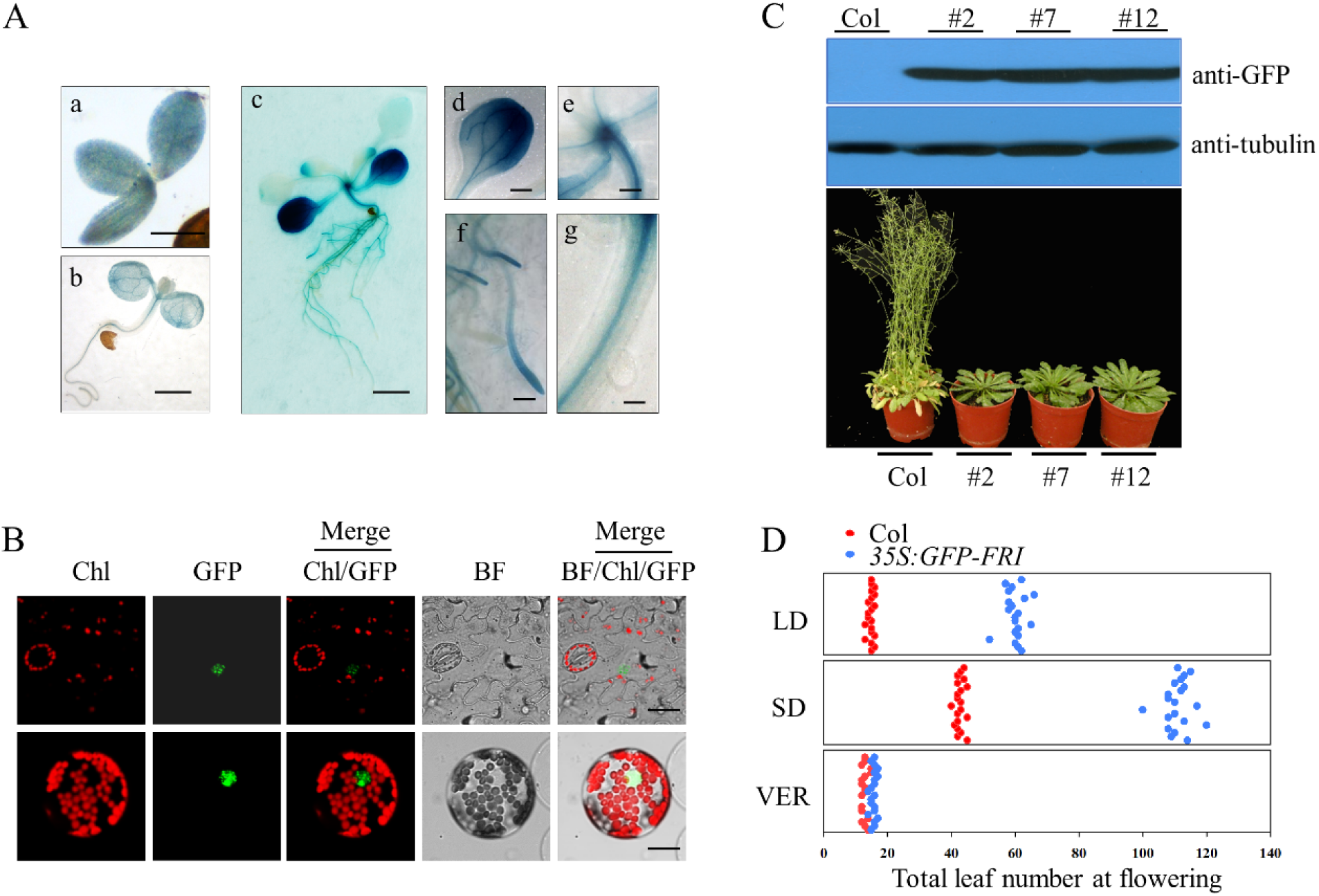
Constitutive expression of *FRI* delays flowering. A. Histochemical analysis of *FRI* expression in the *FRI::GUS*-*FRI* lines. *FRI* expression was indicated by GUS staining of the whole embryo tissue and the seedlings (a, bar=100 μ m; b, bar=0.2 cm; c, bar=0.5 cm). Strong GUS staining was observed in the leaves (d, bar=0.5 cm), meristem (e, bar=0.05 cm), root (f, bar=0.125 cm), and phloem (g, bar=0.01 cm).
B. Localization of *FRI* in plants. The lower leaf epidermis was peeled from transgenic *35S:GFP*-*FRI* plants and GFP fluorescence was observed in the mesophyll cells (upper panel, bar=10 μm). The *35S:FRI*-*GFP* construct was transiently expressed in *Arabidopsis* protoplasts (bottom panel, bar=2 μm).
C. Constitutive expression of *FRI* in the transgenic *35S:GFP*-*FRI* lines resulted in late flowering. Immunoblot analysis of *GFP*-*FRI* expression in the wild-type Col line and three individual transgenic *35S:FRI*-*GFP* lines using anti-GFP antibody (upper panel). The flowering phenotype of Col and three individual lines under long-day conditions (bottom panel).
D. Flowering time as indicated by the total leaf number in plants grown under long-day conditions (LD), short-day conditions (SD) or with 30 d of vernalization treatment at 4°C (VER). The x axis denotes total leaf number. Each dot represents a plant. For each line, 20 plants were scored.

We then generated transgenic line expressing *35S::GFP*-*FRI* and examined *GFP*-*FRI* expression in three individual T3 line. Immunoblot analysis revealed that all of these lines accumulated high levels of *GFP*-*FRI* (Fig. 1C). Consistent with the strong accumulation of *GFP*-*FRI* in these lines, these lines showed delayed flowering compared with wild-type Col in long days (LD, 16-h light/8-h dark) and short days (SD, 8-h light/16-h dark) (Fig. 1C&D, Fig. S1). However, vernalization treatment (30 d of cold) completely abolished the late flowering phenotype of these transgenic lines (Fig. 1D, Fig. S1), similar to findings in the wild-type line. These results confirmed the critical role of *FRI* in delaying flowering time.

### Targeted ectopic expression of *FRI* delays flowering time

*FLC* regulates flowering time in *Arabidopsis* in a spatially-dependent manner, and *FRI* upregulates *FLC* expression (Searle *et al*., 2006). *FRI* itself encodes a large protein (MW >77 KD) that would be incapable of moving long distances, here we also found that *FRI* is widely distributed in different tissue. This possibility prompted us to investigate the spatial effect of *FRI* on flowering time, thus we ectopically expressed *FRI*-*GUS* driven by various tissue-specific promoters, including *pSUC2* and *pRolC* for phloem-specificity, *pKNAT1* for meristem-specificity, *pML1* for leaf epidermis specificity, and *pTobRB7* for root-specificity. GUS staining revealed that these promoters exclusively drove *FRI*-*GUS* expression in specifical tissues (Fig. 2A, Fig. S2). Furthermore, We found that all these transgenic lines of *SUC2::GUS*-*FRI*, *RolC::GUS*-*FRI*, *KNAT1::GUS*-*FRI*, *ML1::GUS*-*FRI* and *TobRB7::GUS*-*FRI* showed delayed flowering (Fig. 2B-C, Fig. S3). Although targeted ectopic expression of *FRI* using spatially specific promoters delayed flowering, the flowering time still varied among these transgenic lines. As shown in Fig. 2B-C, the *SUC2* and *RolC* promoters, which drove *FRI* expression specifically in the phloem, resulted in the greatest delays in flowering, followed by the *KNAT1* promoter and *ML1* promoter. Expressing *FRI* under the control of the *TobRB7* promoter also delayed flowering, but the effect was not as obvious as that of the *SUC2* and *RolC* promoter. Vernalization treatment abolished the late flowering phenotype in these transgenic lines (Fig. S4). These data indicated that *FRI* could function in specific tissues, including the phloem, leaves, shoot meristem and roots, to delay flowering.

**Fig. 2.**
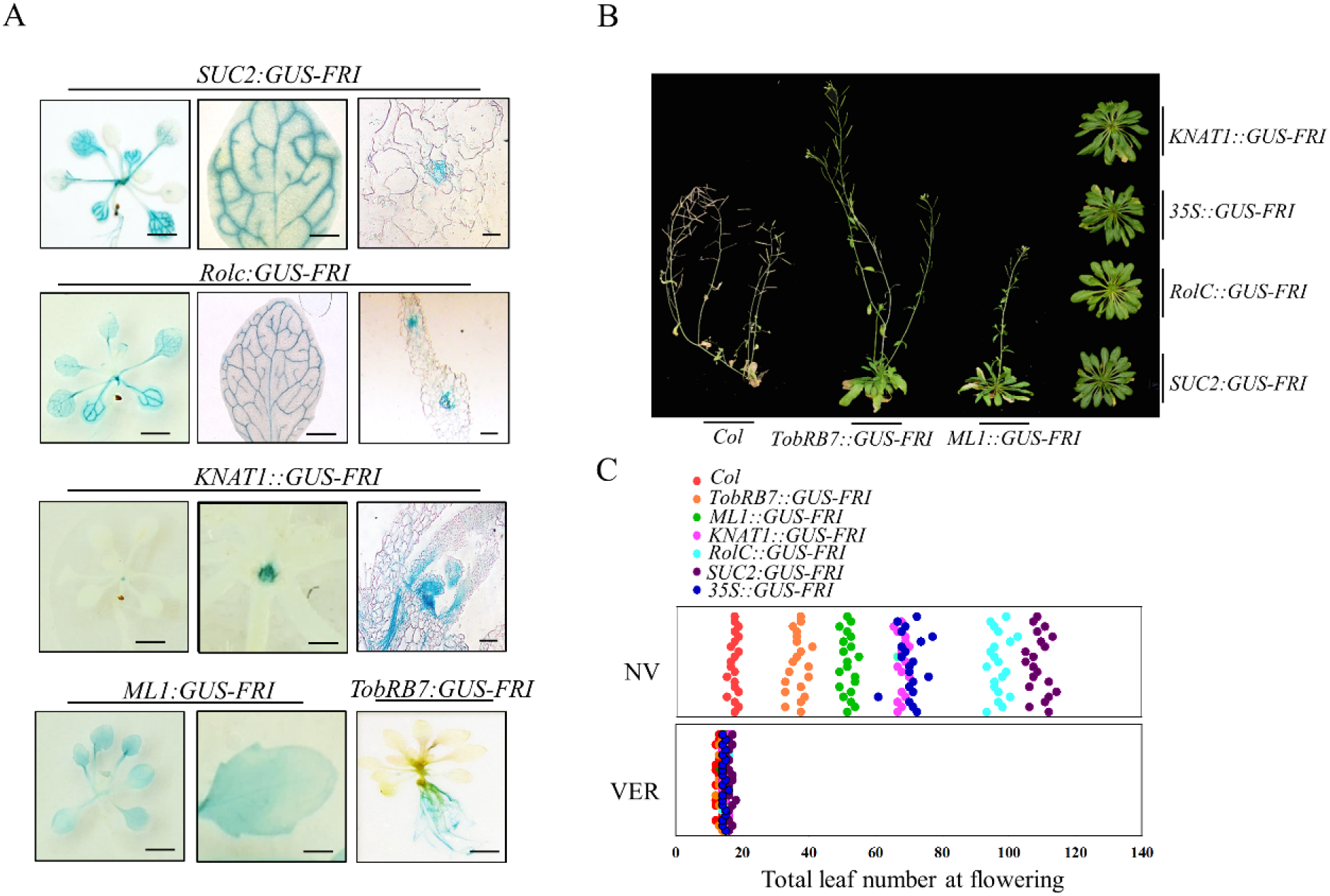
Modulating flowering time by expressing *GUS*-*FRI* in specific tissues. A. Localization of *GUS*-*FRI* expression by GUS staining. GUS staining showed *GUS*-*FRI* expression in the *SUC2::GUS*-*FRI*, *RolC::GUS*-*FRI*, *KNAT1::GUS*-*FRI*, *ML1::GUS*-*FRI* and *TobRB7::GUS*-*FRI* transgenic lines. GUS staining was observed in the whole seedling (left panel; bar=0.5 cm), the leaves (middle panel; bar=0.125 cm), the root (right panel; bar=0.5 cm), or meristem (right panel; bar=0.05 cm). *GUS* staining for the *TobRB7::GUS*-*FRI* line is only presented in the root.
B. Variation in flowering time among various transgenic lines. All of these transgenic lines were grown under long-day conditions, and the flowering phenotypes were presented at 60 days after seed germination.
C. Quantitative flowering time among various transgenic lines as indicated by the total leaf number under long-day conditions without vernalization (NV) or with vernalization treatment at 4°C (VER). The x axis denotes total leaf number. Each dot represents a plant.for each line, 20 plants were scored.

### Spatial expression of *FRI* in the phloem significantly affects *FLC* expression and its histone trimethylation level

The transcriptional levels of *FLC* could be markedly up-regulated by *FRI* before vernalization (Shindo *et al*., 2005; Choi *et al*., 2011). Thus, we examined the patterns of *FRI* and *FLC* expression in these transgenic lines. As shown in Fig. 3A (upper panel), we found that the levels of *FRI* transcript in transgenic lines were consistent with the expression patterns driven by the spatial-specific promoters. The transcriptional level of *FLC* was increased over 30-fold in the leaves of the *SUC2::GUS*-*FRI* and *RolC::GUS*-*FRI* lines in comparison with those of the wild-type Col line. The *FLC* levels in the shoot apices of the *KNAT1::GUS*-*FRI* line and in the root tissue of the *TobRB7::GUS*-*FRI* line were also up-regulated, though to a lesser extent than in the leaves of the *SUC2::GUS*-*FRI* line (Fig. 3A, upper panel). Furthermore, we found that the *FT* transcript level in the leaves and the *SOC1* transcript level in the leaves and shoot apices of all these lines were markedly lower than that of wild-type plants, except the *KNAT1::GUS*-*FRI* line, which had high levels of the *FT* and *SOC1* transcripts in the leaves (Fig. 3A, bottom panel). Vernalization treatment blocked the *FLC* transcription, and increased the transcription of *FT* and *SOC1* in the leaves or shoot apices, in the transgenic lines of *SUC2::GUS*-*FRI*, and also in other transgenic lines (Fig. S5). These data suggest that Spatial expression of *FRI* in the phloem mainly delayed flowering by upregulating the expression of *FLC* in the leaves.

**Fig. 3.**
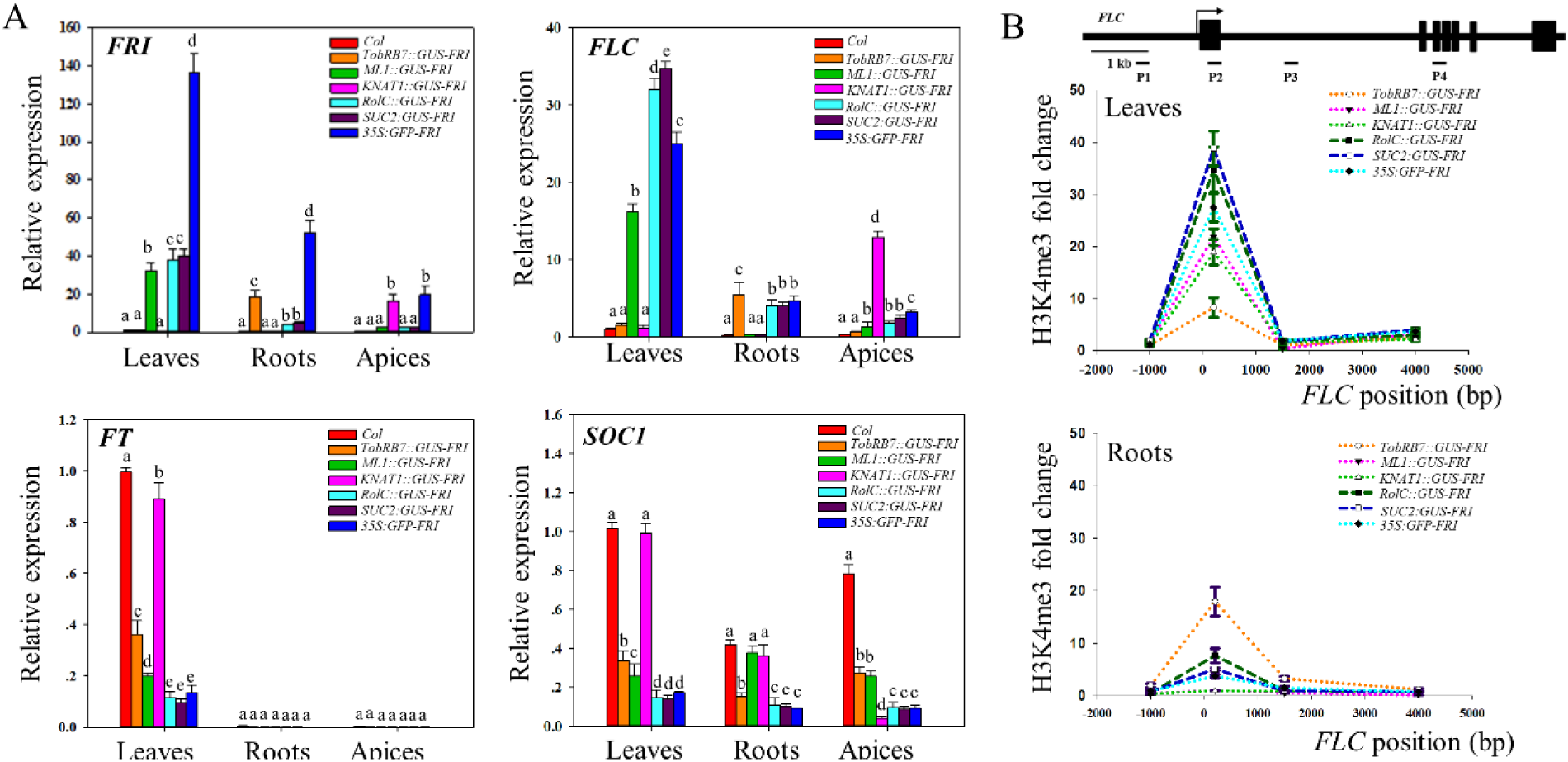
*FRI* effectively promotes the levels of *FLC* transcript and represses *FT* and *SOC1* expression when expressed in the phloem and leaves. A. Quantitative RT-PCR analysis of the spatial expressions of *FRI*, *FLC*, *FT* and *SOC1* in different tissues from different transgenic lines. Ten-day-old seedlings grown under LD conditions. The beta-tubulin gene (*TUB2*) was amplified as an internal control. Values are means ± SD of three biological replicates. Bars with different letters indicate significant differences among different transgenic lines within the same tissue (*p*<0.05, ANOVA followed by Tukey’s post hoc test).
B. CHIP-qPCR analysis of the relative levels of H3K4me3 in *FLC* chromatin. DNA fragments were obtained from the leaves (upper panel) and roots (bottom panel) of ten-day-old plants by CHIP using H3K4me3 antibody, and the amounts of DNA fragments after ChIP were quantified and subsequently normalized to an internal control (*AGAMOUS*). The primer pairs used in the PCR are shown as bars below *FLC*. Exons are shown as black boxes and introns as black lines. Data are means ± SD of triplicate experiments.

*FRI* promotes *FLC* expression by increasing histone methylation at the *FLC* locus, which delays flowering (Song *et al*., 2012). As shown in Fig. 3B, we measured the levels of histone H3Lys4 trimethylation (H3K4me3), the histone mark associated with active *FLC* chromatin in the leaves. Consistent with the high *FLC* transcript pattern in these transgenic lines, the H3K4me3 levels in the *FLC* genomic region were highest in the leaves of the *SUC2::GUS*-*FRI* and *RolC::GUS*-*FRI* lines, followed by the *KNAT1::GUS*-*FRI* and *ML1::GUS*-*FRI* lines. The H3K4me3 levels at the *FLC* locus were very low in the leaves of the *TobRB7::GUS-FRI* line (Fig. 3B). Furthermore, the H3K4me3 levels at the *FLC* locus in the root tissue of the *TobRB7::GUS*-*FRI* line were also up-regulated, though to a lesser extent than in the leaves of the *SUC2::GUS*-*FRI* line. Vernalization treatment completely suppressed the H3K4me3 levels at the *FLC* locus in the leaves and roots of these transgenic lines (Fig. S6).

### Spatial expression of *FRI* in the roots specifically activates *MAF4* and *MAF5* expressions and delays flowering in *FLC*-dependent pathways

A previous study showed that the spatial expression of *FLC* in the root did not delay flowering (Searle *et al*., 2006). In our current experiments, ectopic expression of *FRI* expression in the roots of the *TobRB7::GUS*-*FRI* line delayed flowering, and reduced levels of *FT* and *SOC1* transcripts in the leaves (Fig. 3A), thus it is possible that *FRI* might activate additional genes to delay flowering in the *TobRB7::GUS*-*FRI* line. In addition to *FLC*, there exists five MADS gene (*FLM* and *MAF2*-*5*) homology to *FLC* in *Arabidopsis* genome (Ratcliffe *et al*., 2003; Scortecci *et al*., 2003; Gu *et al*., 2009; Kim *et al*., 2010). Thus, we further compared the transcript differences of *FLM* and *MAF2*-*5* in the roots of *TobRB7::GUS*-*FRI*, *SUC2::GUS*-*FRI*, *35S::GFP*-*FRI* and wild-type lines. We found that there was no obvious change in *MAF1*, *MAF2* or *MAF3* expression in the leaves and root tissue of these transgenic lines (Fig. S7A). However, the *MAF4* and *MAF5* transcript levels in the roots of *TobRB7::GUS*-*FRI* plants were nearly 5-fold greater than in the wild type, whereas they were not increased in the roots of other lines (Fig. 4A). These data indicate that root *MAF4* and *MAF5* are possible other targets of *FRI* for delaying flowering in *Arabidopsis*.

**Fig. 4.**
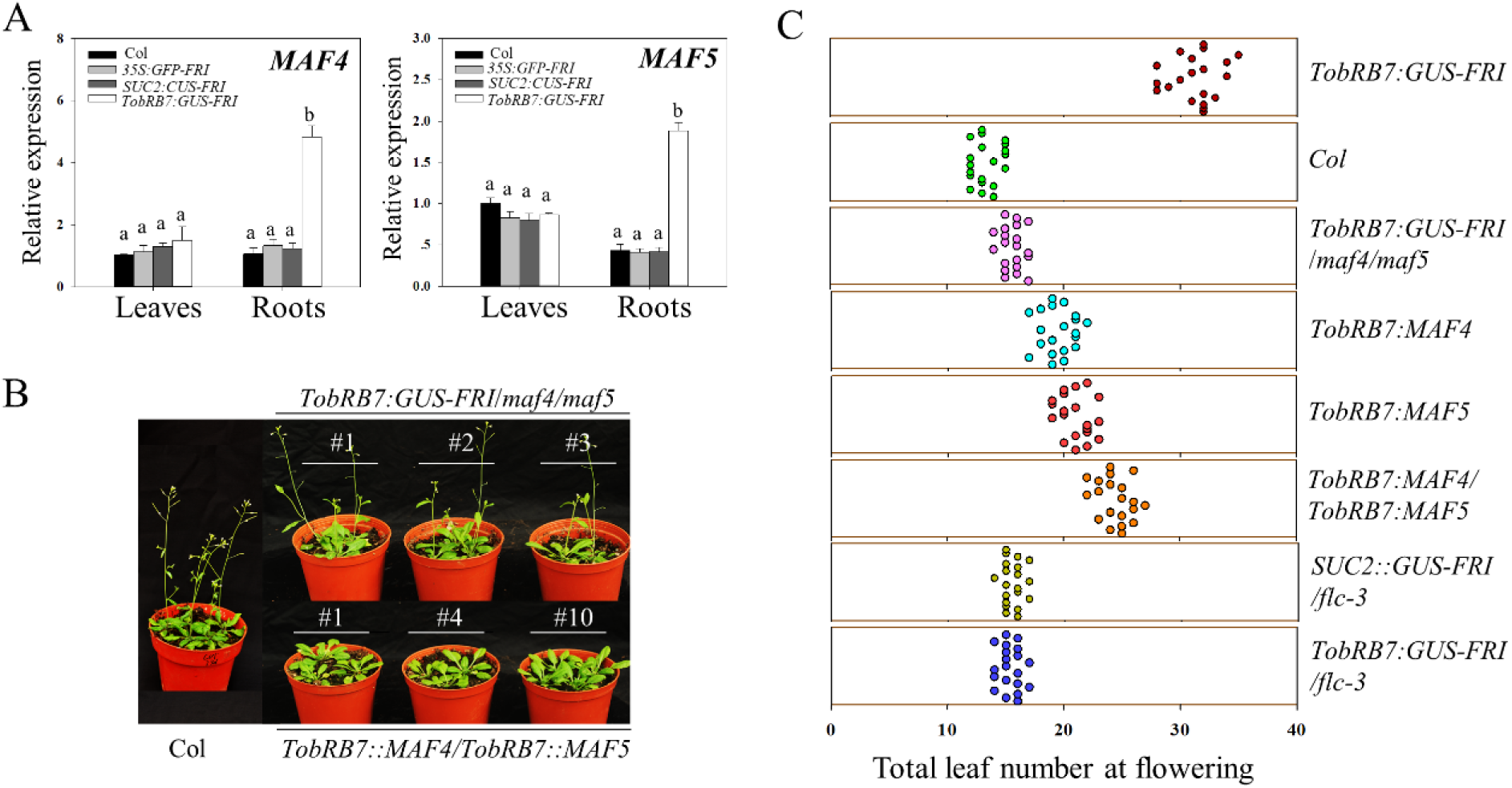
*MAF4* and *MAF5* are specifically activated in the roots of *TobRB7::GUS*-*FRI* line. A. Quantitative RT-PCR analysis of the spatial expressions of *MAF4* and *MAF5* in different tissues from different transgenic lines. Ten-day-old seedlings grown under LD conditions. The beta-tubulin gene (*TUB2*) was amplified as an internal control. Values are means ± SD of three biological replicates. Bars with different letters indicate significant differences among different transgenic lines within the same tissue (*p*<0.05, ANOVA followed by Tukey’s post hoc test).
B. Effects of overexpressing *MAF4 and MAF5* in the root and loss of function of *MAF4 and MAF5* in *TobRB7::GUS*-*FRI* line on flowering time. The plants were grown under long-day (LD) conditions, and the flowering phenotype is presented.
C. Quantitative flowering time among various transgenic lines and mutants as indicated by the total leaf number under long-day conditions. The x axis denotes total leaf number. Each dot represents a plant.for each line, 20 plants were scored.

To further assess the role of root *MAF4* and *MAF5* in controlling flowering time, we obtained two transfer DNA (T-DNA) insertion alleles for *MAF4* and *MAF5* from the *Arabidopsis* Information Resource (TAIR), termed *maf4*-*1* and *maf5*-*1*, respectively. These two alleles failed to produce full-length transcripts, as demonstrated by reverse transcription-PCR (Fig. S7B-C). We then crossed *maf4*-*1* with *maf5*-*1* to obtain the *maf4/maf5*-*1* double mutant (Fig. S7B), and introduced *TobRB7::GUS*-*FRI* into the *maf4/maf5* double mutant (termed *TobRB7::GUS-FRI/maf4/maf5*), and then compared the flowering times with *TobRB7::GUS*-*FRI* line (Fig. 4B-C). Under long-day conditions, knocking out *MAF4* and *MAF5* in the *TobRB7::GUS*-*FRI/maf4/maf5* line dramatically reduced flowering time, in contrast to *TobRB7::GUS*-*FRI*, suggesting that *MAF4* and *MAF5* are required for the late flowering phenotype of *TobRB7::GUS*-*FRI* plants. We then generated transgenic *TobRB7::MAF4* line and *TobRB7::MAF5* line in which *MAF4* or *MAF5* expression was respectively driven by the root-specific *TobRB7* promoter and crossed the *TobRB7::MAF4* line with the *TobRB7::MAF5* line (referred to as *TobRB7::MAF4/TobRB7::MAF5*). As shown in Fig. 4B-C and Fig. S8, in contrast to the wild type, these transgenic plants had late flowering phenotypes, and the double overexpression plants (*TobRB7::MAF4/TobRB7::MAF5*) flowered later than the *TobRB7::MAF4* or *TobRB7::MAF5* lines alone, suggesting that *MAF4 and MAF5* delay flowering when they are expressed specifically in the root.

To confirm whether the late flowering phenotype in the *TobRB7::GUS*-*FRI* line depends on *FLC*, we crossed *SUC2::GUS*-*FRI* or *TobRB7::GUS*-*FRI* with *flc*-*3* (referred to as *SUC2::GUS*-*FRI/flc*-*3* and *TobRB7::GUS*-*FRI/flc*-*3* respectively) and then compared the flowering time of these lines. As previously reported (Searle *et al*., 2006), the loss-of-function *flc*-*3* line showed early flowering under LD or SD conditions (Fig. 9A-B). Under long-day conditions, knocking out *FLC* in the *SUC2::GUS*-*FRI/flc*-*3* line markedly promoted flowering, resulting in a flowering time that matched that of the wild type (Fig. 4C), suggesting that *FLC* was mainly responsible for the late flowering time of the *SUC2::GUS*-*FRI* line. In addition, mutation of *FLC* in the *TobRB7::GUS*-*FRI/flc*-*3* background also dramatically promoted flowering in comparison with the *TobRB7::GUS*-*FRI* line, suggesting the synergistic effect of *FLC* and *MAF4*/*MAF5* in regulating flowering time in *TobRB7::GUS*-*FRI* line.

### Root *MAF4* and *MAF5* mediate *FRI*-dependent late flowering through antagonizing leaves *FT* signal

Grafting is a useful tool for investigating the movement of long-range signals within a plant. For instance, grafting experiments demonstrated that FT moves from leaves to the shoot meristem to initiate flowering (An *et al*., 2004). To evaluate whether certain mobile signals exist that travel from the roots to shoot to delay flowering once *FRI* is expressed in the root tissue, we grafted the rootstock of *TobRB7::GUS*-*FRI* with wild-type Col scion (termed *TobRB7::GUS*→*FRI* Col). Such grafted line had a later flowering time than the control line, in which wild-type Col rootstock was grafted to wild-type Col scion (termed Col→Col) (Fig. 5A-B, Fig. 9C-D). The *FT* level in the leaves of *TobRB7:: GUS*-*FRI*→ Col line was lower than that in the control Col→Col, and the expressions of *MAF4* and *MAF5* were dramatically increased in the roots of *TobRB7::GUS*-*FRI*→ Col plants (Fig. 5C). Similarly, in contrast to the grafted Col→ Col line, the graft *TobRB7::MAF4/TobRB7::MAF5*→Col line also exhibited delayed flowering (Fig. 5A-B, Fig. 9C-D). These data indicated that a potential mobile factor travels from the rootstock of *TobRB7::GUS*-*FRI* or *TobRB7::MAF4/TobRB7::MAF5* to suppress floral initiation in the wild-type Col scion, and such mobile factor possibly interferes with the florigen *FT* signal to reduce *FT* expression in the leaves.

**Fig. 5.**
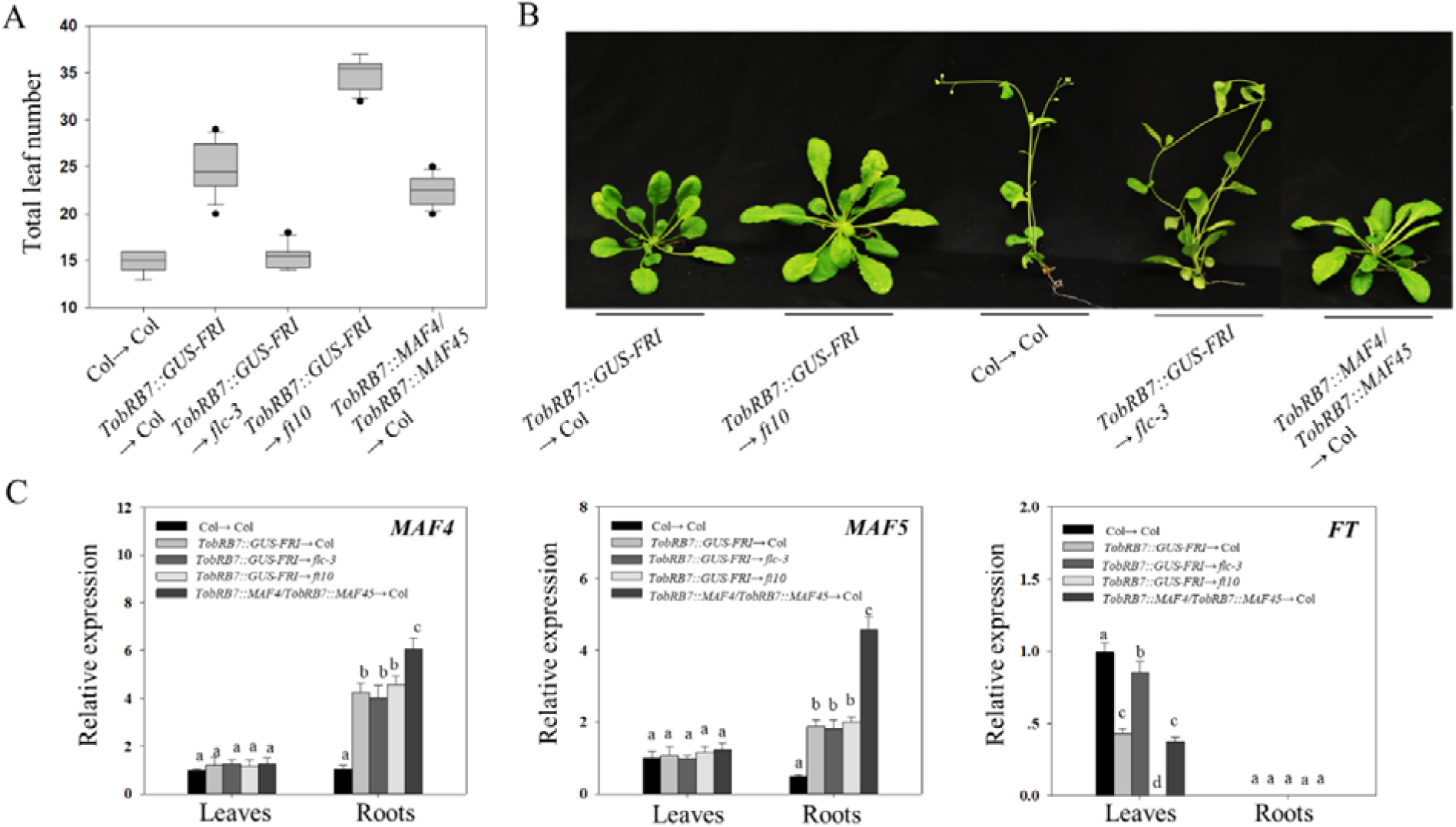
Variation in flowering time among different grafts with *TobRB7::GUS*-*FRI* as the rootstock. (A) Flowering time and phenotypes (B) among different grafts with *TobRB7::GUS*-*FRI* or *TobRB7::MAF4/ TobRB7::MAF5* as the rootstock. The flowering time under LD conditions is given as the total leaf number at the time of flowering, and the flowering phenotypes were presented at 40 days after seed germination. Data are means ± SD of three replicates (n=12). (C) Quantitative RT-PCR analysis of the expressions of *MAF4*, *MAF5* and *FT* in different tissues from different grafts. Ten-day-old seedlings grown under LD conditions. The beta-tubulin gene (*TUB2*) was amplified as an internal control. Values are means ± SD of three biological replicates. Bars with different letters indicate significant differences among different transgenic lines within the same tissue (*p*<0.05, ANOVA followed by Tukey’s post hoc test).

To further test the role of *FT* in mediating the late flowering phenotype of the *TobRB7::GUS*-*FRI* line, we introduced *TobRB7::GUS*-*FRI* into the late flowering *ft*-*10* mutant (termed *TobRB7::GUS*-*FRI/ft*-*10*). As expected, flowering was markedly delayed in *TobRB7::GUS*-*FRI/ft*-*10* plants, as in the *ft*-*10* mutant (Fig. 9A-B). The grafted *TobRB7::GUS*-*FRI*→*ft*-*10* line(*TobRB7::GUS*-*FRI* rootstock connected to *ft*-*10* scion) showed a similar late flowering time as the *ft*-*10* mutant, suggesting the late flowering phenotype of the *TobRB7::GUS*-*FRI* line requires the leaf *FT* signal (Fig. 5 A-B, Fig. 9C-D). Furthermore, we found that the *TobRB7::GUS*-*FRI*→*flc*-*3* line (*TobRB7::GUS*-*FRI* rootstock to the *flc*-*3* scion) flowered earlier than did the *TobRB7::GUS*-*FRI*→Col line (Fig. 5A-B, Fig. 8C-D). These data further support that a mobile signal that functions as an anti-florigen travels from the rootstock of *TobRB7::GUS*-*FRI* to delay flowering, and the presence of functional *FLC* in the leaves has a certain function in the late flowering phenotype of *TobRB7::GUS*-*FRI*.

### Misexpressing *FRI* in the embryo delays flowering time and increases H3K4me3 level at *FLC* locus

Beside the spatial effect of *FRI* on flowering time, we also investigate the temporal effect of *FRI* on flowering, because epigenetic modification, which is characteristic of long-term temporal effect, play the essential role in *FRI*-dependent flowering time (Song *et al*., 2012; Wigge *et al*., 2005). To achieve it, we investigated the effect of *FRI* expression at different developmental stages. We firstly generated transgenic lines in which the expression of *GUS*-*FRI* fusions was driven by two embryo-specific promoters: *LEC2*, for early embryogenesis specificity, and *EM1*, for seed maturation expression (Braybrook and Harada, 2008; Gaubicr *et al*., 1993). As shown in Fig. 6A, GUS staining was particularly strong in the embryos of these transgenic lines, suggesting that the *EM1* and *LEC2* promoters are specifically activated *GUS*-*FRI* in the embryonic tissue (Fig. 6A). We found that all of these lines had a delayed-flowering phenotype in LD conditions (Fig. 6B-C). These transgenic lines also had high levels of *FLC* expression and H3K4me3 modification within the *FLC* locus in the embryo compared with the wild type (Fig. 6D-E), suggesting that *FRI* expression at the embryonic stage efficiently delays flowering time, possibly through epigenetic modification of the *FLC* locus. We also found that vernalization treatment abolished H3K4me3 modification at the *FLC* locus in these transgenic lines to promote flowering, similar to the wild type (Fig. 6C-E, Fig. S10).

**Fig. 6.**
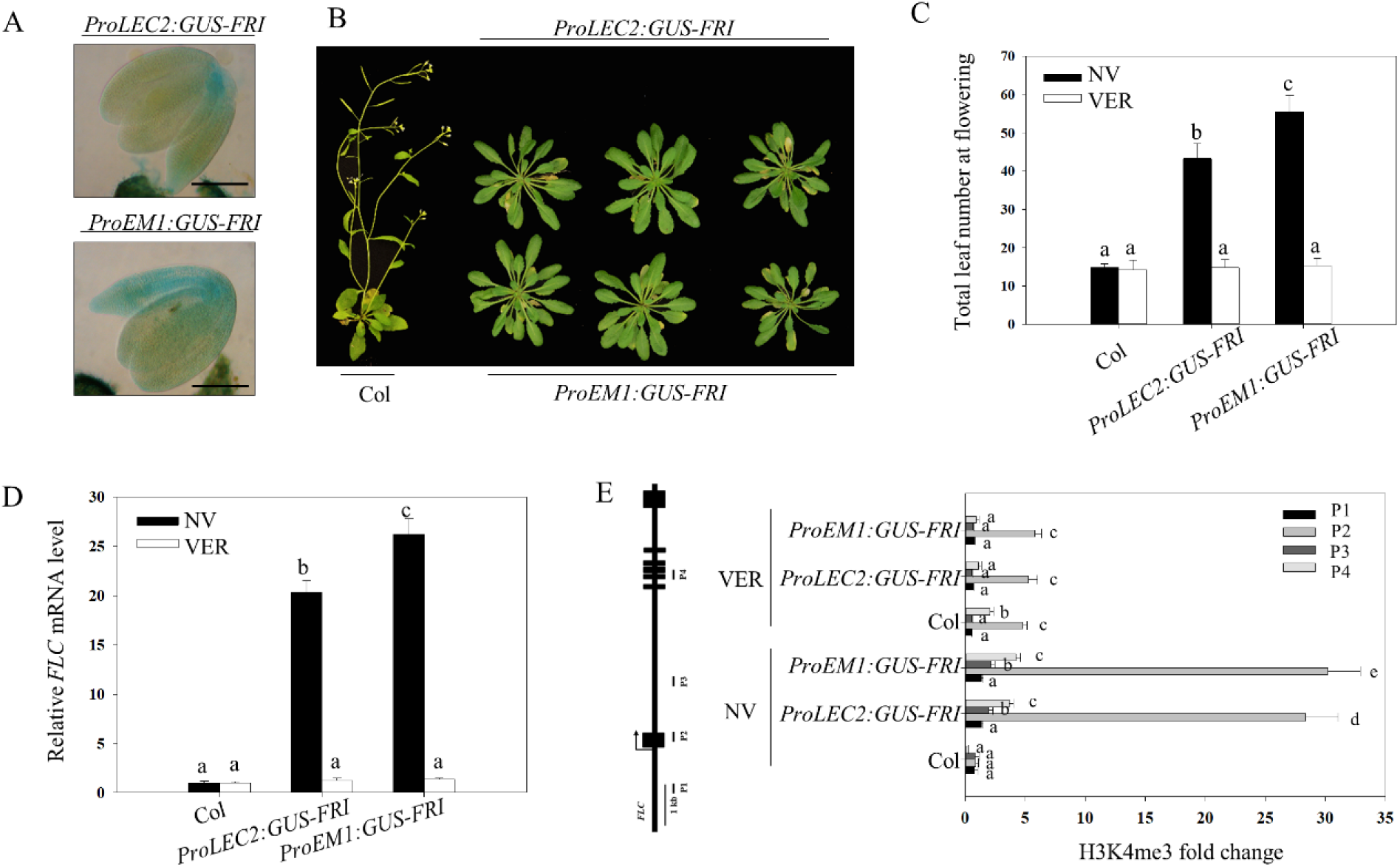
Modulating flowering time by expressing *GUS*-*FRI* driven by seed-specific promoters. A. Localization of *GUS*-*FRI* expression by GUS staining. GUS staining showed that *FRI* is specifically expressed in the seed of *ProLEC2::GUS*-*FRI* and *ProEM1::GUS*-*FRI* transgenic lines (bar=100 μ m).
B. Phenotypes and (C) flowering time of *ProLEC2::GUS*-*FRI* and *ProEM1::GUS*-*FRI* transgenic lines. The flowering phenotypes were presented at 40 days after seed germination. The flowering time is given as the total leaf number at the time of flowering under long-day conditions without vernalization (NV) or with vernalization treatment (VER). Data are means ± SD of three replicates. For each line, 20 plants were scored. Bars with different letters are significantly different at *p*<0.05.
C. Quantitative RT-PCR analysis of the expressions of *FLC* in different transgenic lines. Ten-day-old seedlings grown under LD conditions without vernalization (NV) or with 30 d of vernalization treatment at 4°C (VER). The beta-tubulin gene (*TUB2*) was amplified as an internal control. Values are means ± SD of three biological replicates. Bars with different letters indicate significant differences among different transgenic lines(*p*<0.05).
D. CHIP-qPCR analysis of the relative levels of H3K4me3 in *FLC* chromatin. DNA fragments were obtained from ten-day-old seedlings with or without vernalization (VER or NV) by CHIP using H3K4me3 antibody, and the amounts of DNA fragments after ChIP were quantified and subsequently normalized to an internal control (*AGAMOUS*). The primer pairs used in the PCR are shown as bars below *FLC*. Exons are shown as black boxes and introns as black lines. Data are means ± SD of triplicate experiments.

### Misexpressing *FRI* at different developmental stages affect the flowering time and H3K4me3 level at *FLC* locus

To determine which development status is prerequisite for *FRI*-dependent flowering time, we applied the estrogen-inducible system by driven *FRI* with the estrogen-inducible promoter (Zuo *et al*., 2000). We inserted the *GUS*-*FRI* fusion downstream of the lexA-binding domain, and transgenically expressed this construct (*ProER8:GUS*-*FRI*) in *Arabidopsis* by *Agrobacterium*-mediated transformation. As shown in Fig. S11, the GUS reporter gene was induced after 24 h of estradiol treatment in the transgenic *ProER8:GUS-FRI* line. We then treated the transgenic *ProER8:GUS*-*FRI* line with estradiol at different development period to see the flowering variance. We firstly tested the induction of *FRI* expression in the pro-embryo to post-embryonic period. Exogenous application of estradiol in the flower bud formation stage (2 days before self-pollination) markedly delayed flowering in the first generation (Fig. 7A-C). However, estradiol treatment for the flowers at the day, 1 day, 3 days, 5 days or 7 days after artificial pollination had different effects on flowering of the next generation. As shown in Fig. 7A-C, estradiol treatment at the day and one day after pollination, the progeny plants flowered very late. In contrast to the late flowering phenotype, estradiol treatment at 3 days after pollination had weak effect on flowering of the next generation, especially in the later seed formation stages. The levels of *FLC* transcript and the H3K4me3 levels at the *FLC* locus in the progeny plants were coincide with the flowering time under LD conditions (Fig. 7D-E). These data suggest that *FRI* expression in the pro-embryo to post-embryonic period affects flowering time, and that estradiol induced *FRI* expression during gametogenesis and early embryogenesis has programed epigenetic modifications in the *FLC* chromatin of the next generation, but estradiol appeared not to penetrate the developing seed coat to influence flowering of the progeny plants when application of estradiol at later embryogenesis.

**Fig. 7.**
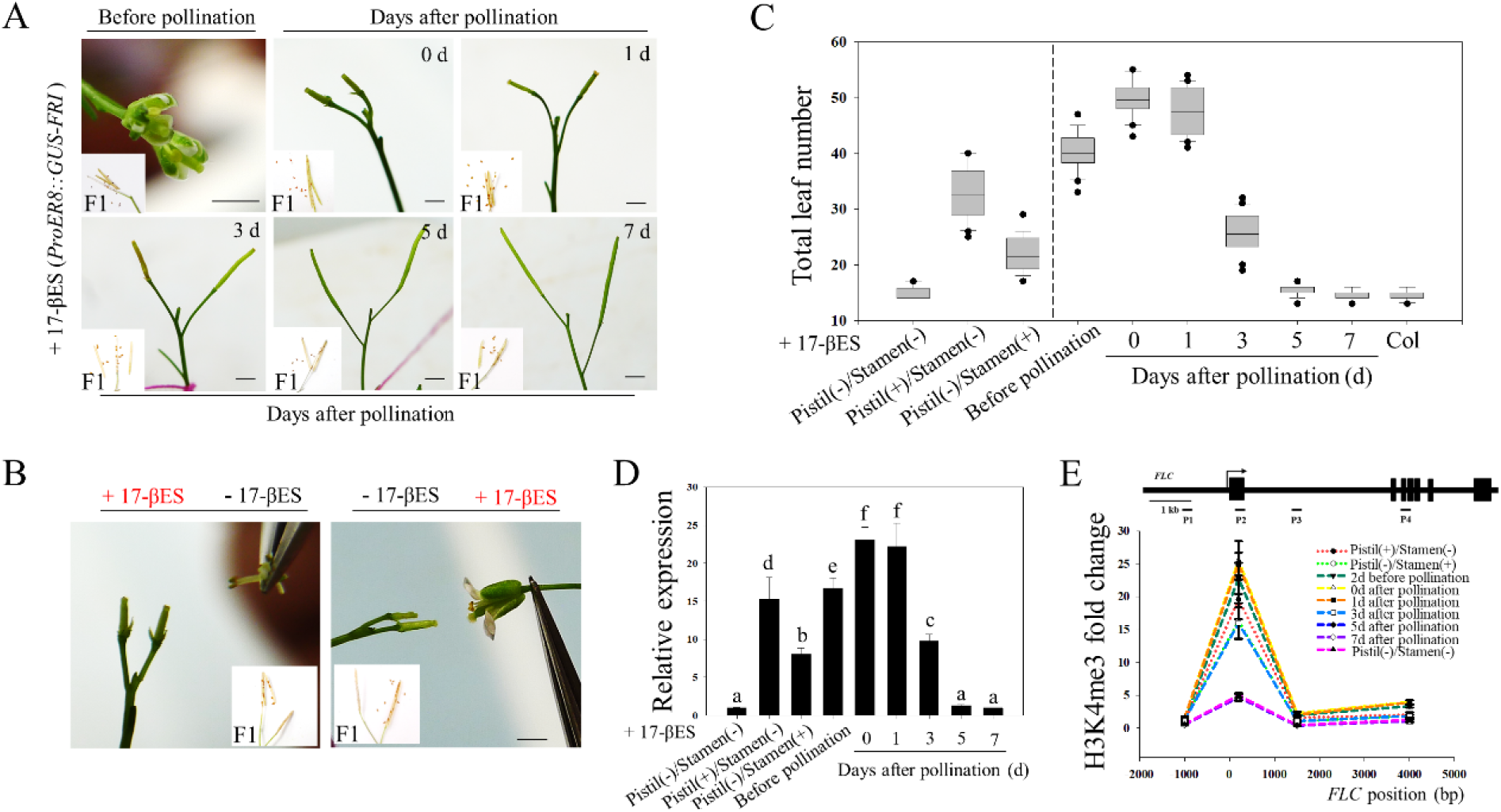
Modulating flowering time by expressing *GUS*-*FRI* in inducible *ProER8:GUS*-*FRI* line during pro-embryo to post-embryonic stage. A. Estradiol induction in different embryonic development stages of *ProER8:GUS*-*FRI* line. Estradiol was applied to parent plant during reproductive development, from flower to seed formation. Seeds were harvested from mother plants after maturity. bar=2 mm.
B. Estradiol induction in pistil or Stamen of *ProER8:GUS*-*FRI* line before pollination. Pistil with estradiol-treated was crossed with stamen without estradiol-treated, or Pistil without estradiol-treated was crossed with stamen with estradiol-treated. Seeds were harvested from mother plants after maturity. bar=2 mm.
C. Flowering time of progeny after parent plants being treated with estradiol at the pro-embryo to post-embryonic stage. The flowering time under LD conditions is given as the total leaf number at the time of flowering. Data are means ± SD of three replicates. For each line, 20 plants were scored.
D. Quantitative RT-PCR analysis of the expressions of *FLC* in the progeny. Ten-day-old seedlings grown under LD conditions. The expressions of *FLC* were measured in the leaves of different progeny plants. The beta-tubulin gene (*TUB2*) was amplified as an internal control. Values are means ± SD of three biological replicates. Bars with different letters indicate significant differences among progeny plants after different treatments to the parent plants (*p*<0.05).
E. CHIP-qPCR analysis of the relative levels of H3K4me3 in *FLC* chromatin of transgenic *ProER8:GUS*-*FRI* line among different progeny plants. Ten-day-old seedlings grown under LD conditions were obtained by CHIP using H3K4me3 antibody, and the amounts of DNA fragments after ChIP were quantified and subsequently normalized to an internal control (*AGAMOUS*). The primer pairs used in the PCR are shown as bars below *FLC*. Exons are shown as blue boxes and introns as black lines. Data are means ± SD of triplicate experiments.

To further assess the contribution of *FRI* in maternal and paternal gametes to controlling flowering time, we treated flower buds (2 days before self-pollination) with estradiol and crossed the female parent that with or without estradiol treatment with the male parent that without or with estradiol treatment by emasculation and artificial pollination. We found that the progeny by crossing the pistil (with estradiol treatment) with the stamen (without estradiol treatment) exhibited delayed flowering. The progeny by crossing the pistil (without estradiol treatment) with the stamen (with estradiol treatment) also delayed flowering, though the effect of delaying flowering was greatly reduced (Fig. 7B). In addition, the *FLC* expression levels and the H3K4me3 levels at the *FLC* locus of the progeny by crossing the pistil (with estradiol treatment) with the stamen (without estradiol treatment) were higher than that of the progeny by reverse crossing (Fig. 7D-E). Our findings suggest that estradiol induced *FRI* expression during female gametogenesis is more inheritable and stable than that during male gametogenesis through epigenetic modifications in the *FLC* chromatin of the next generation.

Furthermore, estradiol treatment at the seed stage, 5 days and 7 days old seedling stages markedly delayed flowering. However, the effect of inhibiting flowering was greatly reduced when the 9 days old seedlings were treated with estradiol. Moreover, 11 days, 13 days or 15 days old seedlings did not delay flowering after estradiol treatment (Fig. 8A-B). These data indicate that flowering is only delayed if *FRI* is expressed in the embryo or in young seedlings. We then measured *FRI*, *FLC*, *FT*, and *SOC1* expression and H3K4me3 levels at the *FLC* locus in the *ProER8:GUS*-*FRI* line treated with estradiol at different growth stage. As shown in Fig. 8C and Fig. S12, the *FRI* expression was induced after 12 h of estradiol treatment in all transgenic *ProER8:GUS*-*FRI* lines of different stages. The levels of *FLC* transcript were high and those of *FT* and *SOC1* transcript were low in the induced seedlings. However, the *FLC* expression levels were reduced and the *FT* and *SOC1* transcript levels were increaced when the 9-15 days old seedlings treating with estradiol, compared with the younger seedlings. The H3K4me3 levels and H3K27me3 levels at the *FLC* locus were coincide with the expression pattern of *FLC* (Fig. 8D). Recently, a new study found that FT can suppress *FLC* mRNA expression and change the *FLC* transcription states (Chen and Penfield, 2018). In this study, the lower levels of *FLC* transcript and higher levels of H3K27me3 in *FLC* chromatin in older seedlings is probably due to the regulation of FT protein existed in the leaves. Consistent with this, even though the *FT* and *SOC1* mRNA expression were suppressed by *FLC* in older seedlings, but the plants showed a similar early flowering time as the non-induced seedlings. Together, these data indicate that *FRI* must be expressed at early development stage to ensure the epigenetic activation of *FLC* expression for late flowering.

**Fig. 8.**
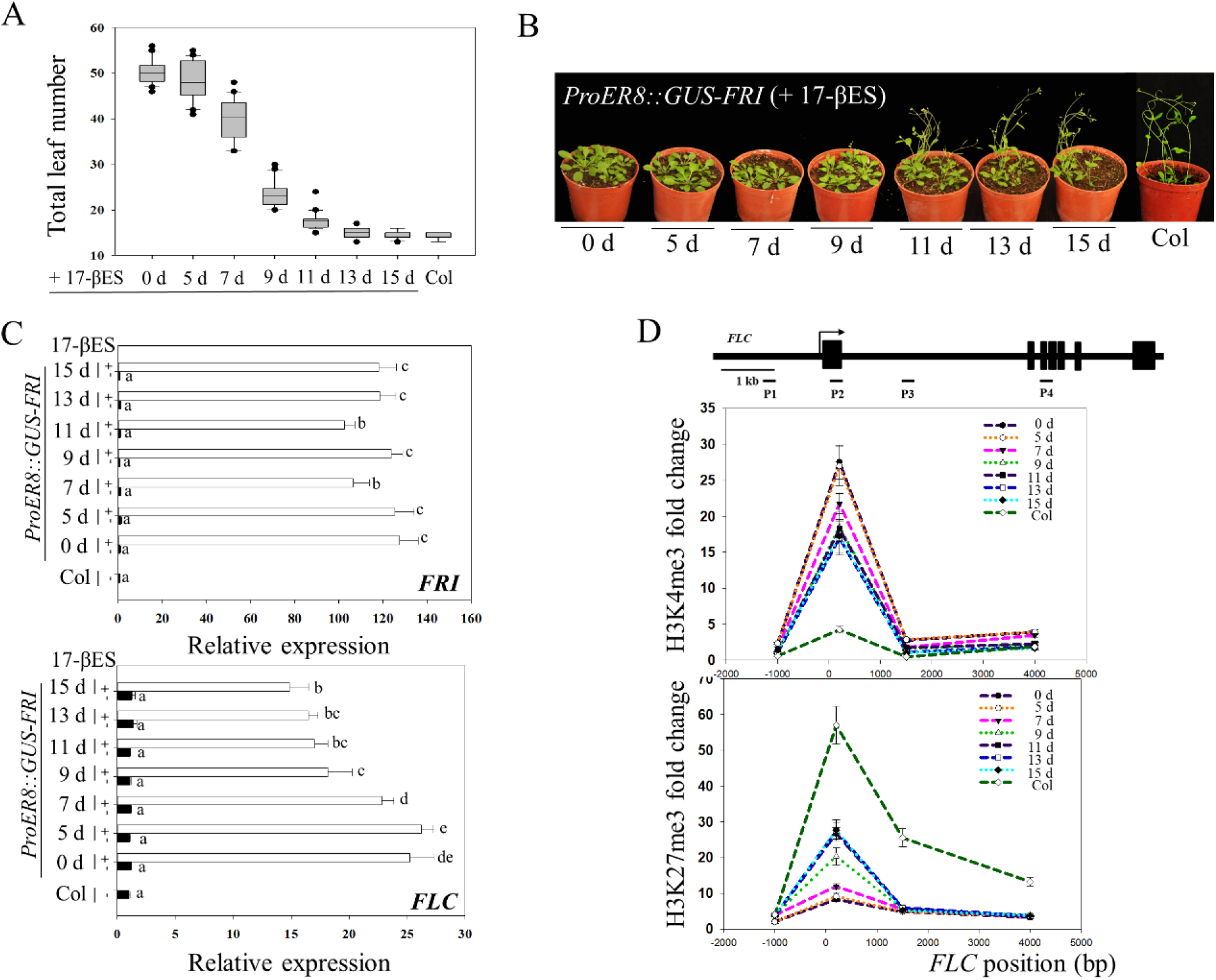
Modulating flowering time by expressing *GUS*-*FRI* in inducible *ProER8:GUS*-*FRI* line in young seedlings. (A)Flowering time and (B) phenotypes of transgenic *ProER8:GUS*-*FRI* line treated with estradiol at the embryonic stage or 5, 7, 9, 11, 13, 15 days after germination. The flowering time under LD conditions is given as the total leaf number at the time of flowering, and the flowering phenotypes were presented at 40 days after seed germination. Data are means ± SD of three replicates. For each line, 20 plants were scored. (C)Quantitative RT-PCR analysis of the expressions of *FRI* and *FLC* in the *ProER8:GUS*-*FRI* line treated with or without estradiol at the embryonic stage or 5, 7, 9, 11, 13, 15 days after germination. The seeds or seedlings grown under LD conditions were harvested 2 days after treated with or without estradiol. The beta-tubulin gene (*TUB2*) was amplified as an internal control. Values are means ± SD of three biological replicates. Bars with different letters indicate significant differences among different treatments (*p*<0.05). (D)CHIP-qPCR analysis of the relative levels of H3K4me3 and H3K27me3 in *FLC* chromatin of transgenic *ProER8:GUS*-*FRI* line treated with estradiol. The seeds or seedlings grown under LD conditions were harvested 5 days after treated with or without estradiol. DNA fragments were obtained from the seeds or seedlings by CHIP using H3K4me3 or H3K27me3 antibody, and the amounts of DNA fragments after ChIP were quantified and subsequently normalized to an internal control (*AGAMOUS*). The primer pairs used in the PCR are shown as bars below *FLC*. Exons are shown as black boxes and introns as black lines. Data are means ± SD of triplicate experiments.

## Discussion

### *FRI* targets root *MAF4* and *MAF5* to regulate flowering time

For most plants, day length and environmental temperature are important flowering cues (Jaeger *et al*., 2007; Amasino and Michaels, 2010). Since flower development occurs at the shoot apical meristem or in the lateral meristem, but photoperiod and ambient temperature signals are perceived by the growing leaves, it is not surprising that the flowering signal is communicated among spatially separated organs in plant. *FT* acts both in the phloem and meristem to trigger flowering (Corbesier *et al*., 2007; Wigge, 2011). Misexpression of *FLC* from a phloem-specific promoter also represses flowering by reducing *FT* transcript levels in the leaves, and the late flowering phenotype of such plant can be overcome by expressing *FT* in the same tissue (Searle *et al*., 2006). In our experiments, we found that *FRI* is globally expressed in different tissues, including leaves, meristems, phloem, and roots. The spatial expression of *FRI* in the phloem prominently activated *FLC* expression and delayed flowering time. Furthermore, Our results showed that ectopic expression of *FRI* in the root tissue of *TobRB7::GUS*-*FRI* plants also repressed flowering, though these plants flowered earlier than did the *SUC2::GUS*-*FRI* line.

Previous studies mainly focused on the role of leaves in perceiving light or ambient signals to initiate flowering (An *et al*, 2004; Searle *et al*., 2006, Amasino and Michaels, 2010). However, in *Sinapis alba*, sucrose triggers the release of cytokinins from the roots during photoperiodic treatment, which is necessary for floral induction (Bernier *et al*., 1993). Recently, global transcriptome analysis identified 595 genes, including 18 known flowering time genes, which were differentially expressed in the root tissue under inductive long-day conditions (Bouche *et al*, 2016), suggesting that root tissue is integrated into the whole plant network governing flowering time. A previous study showed that ectopic expression of *FLC* in the root did not delay flowering (Searle *et al*., 2006), implying that *FRI* upregulates new target to delay flowering in the *TobRB7::GUS*-*FRI* line. It has been reported that *FLC* homologues, *FLOWERING LOCUS M (FLM)/MADS AFFECTING FLOWERING 1*, and *MAF2*-*MAF5*, are also associated with flowering time (Ratcliffe *et al*., 2003; Scortecci *et al*., 2003; Gu *et al*., 2009; Kim *et al*., 2010). Here we excluded *MAF1*-*MAF3* as the potential targets of *FRI* in the *TobRB7::GUS*-*FRI* line, because their expressions were not markedly upregulated in the *TobRB7::GUS*-*FRI* line. However, two other possible targets, *MAF4* and *MAF5*, were specifically upregulated in the roots of the *TobRB7::GUS*-*FRI* line, but not in the roots and leaves of *SUC2::GUS*-*FRI* plants, suggesting that root *MAF4* and *MAF5* in the *TobRB7::GUS*-*FRI* line may function in mediating flowering.

We then observed that overexpressing *MAF4 and MAF5* in the root indeed delayed flowering than the wild type, whereas mutation of *MAF4 and MAF5* in the *TobRB7::GUS*-*FRI/maf4/maf5* line promoted flowering, resulting in early flowering than in the *TobRB7::GUS*-*FRI* plants. These data support the notion that *FRI* targets *MAF4* and *MAF5* to delay flowering in the *TobRB7::GUS*-*FRI* line. Previous studies also reported that *FLOWERING LOCUS C* clade members act in partial redundancy in floral repression and mediate flowering responses (Gu *et al*., 2013). Here we found that mutation of *FLC* in the *TobRB7::GUS*-*FRI* line markedly promoted flowering, suggesting that *FLC* is also required for the late-flowering phenotype of the *TobRB7::GUS*-*FRI* line, possibly via combined function of *FLC* and *MAF4/MAF5* in regulating flowering.

### Spatial expression of *FRI* or *MAF4/MAF5* in the root antagonizes the *FT* signal in leaves to delay flowering

In our experiments, the transgenic expression of *FRI* in the root activated *MAF4* and *MAF5* expressions and repressed the expression of *FT* in the leaves, and its downstream target *SOC1* in the shoot apex, thereby delaying flowering. These data suggest that the *FT* signal functions in the late flowering process by modulating the spatial expression of *FRI* or *MAF4/MAF5* in the root. Furthermore, our grafting experiments showed that, as in *TobRB7::GUS*-*FRI* plants, the *FT* transcript levels in the scion leaves of *TobRB7::GUS*-*FRI*→ Col was low compared to the wild-type Col. Nevertheless, we observed that the *TobRB7::GUS*-*FRI/ft*-*10* plants and the grafted *TobRB7:: GUS-FRI*→*ft*-*10* plants had similar flowering times as the *ft*-*10* mutant. These data suggest that systemic signals may exist that move from the rootstock to the shoot, where they possibly act downstream of *FRI* to antagonize *FT* expression to delay flowering. We speculate that this systemic signal may either be antiflorigen or a floral repressor. Antiflorigen was previously proposed to oppose florigen and was identified as a mobile signal that precisely controlled flowering time (Matsoukas, 2015). In *Arabidopsis*, *RELATIVE OF CENTRORADIALIS* (*ATC*) is regarded as antiflorigen, since *FT* and *ATC* both interact with *FD* to affect the same downstream genes, such as *AP1*, but have opposite effects on their expression (Huang *et al*., 2012). Small non-coding RNAs, such as *miR156* or *miR172*, may also function as antiflorigen to regulate the juvenile-to-adult and vegetative-to-reproductive phase transitions in several plant species (Wang *et al*., 2009; Wu *et al*., 2009, Skopelitis *et al*., 2018). Transcriptional and posttranscriptional gene silencing that are mediated by mobile signal can travel long distance from the root to the shoot through phloem (Liang *et al*., 2012). Deng *et al*. reported that an inverted-repeat RNA targets intronic regions to promote *FT* expression in *Arabidopsis* (Deng and Chua, 2015). Therefore, it is possible that the systemic signals that travel from the roots of *TobRB7:GUS*-*FRI* plants to the shoot apices are small mobile proteins or non-coding RNAs that inhibit flowering. In agreement with this, long-distance GFP silencing signal can travel from the root to the shoot, and then suppresses GFP expression in the 35S-GFP scion leaves (Liang *et al*., 2012). Thus, more work is needed to identify this signal and understand how it cooperates with *FT* to accurately modulate flowering time.

### Expressing *FRI* during embryonic development and in the young seedling stage efficiently delayed flowering time

It is reported that the seed-specific transcription factor *LEAFY COTYLEDON1* (*LEC1*) promotes the initial establishment of an active chromatin state at *FRI*-mediated *FLC* and activates its expression de novo in the pro-embryo (Tao *et al*., 2017). Both of *LEC1* and *LEC2* are master regulators for seeds development, *LEC2* directly interacts with *LEC1* (NF-YB9) in developing seeds, and *EM1* is specifically expressed in the embryo (Boulard *et al*., 2018, Gaubicr *et al*., 1993). To investigate the temporal effect, in particular, during gametogenesis, pro-embryogenesis and post-embryogenesis, on *FRI*-mediates flowering time, we used an estradiol-inducible expression system, or mis-expressed *FRI* using embryo-specific promoters. We found that *FRI* delays flowering when expressed using seed-specific promoters, and that this was accompanied by high H3K4me3 levels at the *FLC* locus and high levels of *FLC* expression. Furthermore, estradiol induced *FRI* expression during gametogenesis and early embryogenesis delayed flowering by epigenetic modifications in the *FLC* chromatin of the next generation. These results are consistent with previous study which showed that the epigenetic reprogramming of chromatin state occurred as early as the time of gametogenesis (Tao *et al*., 2017). Activation of *FRI* during female gametogenesis had more contribution to delaying flowering than activation of *FRI* during male gametogenesis, suggesting that matrilineal inheritance is more important than paternal inheritance in embryonic epigenetic programming at the *FLC* locus.

We also found that estradiol treatment at the embryo stage or small seedlings of less than one-week-old markedly delayed flowering, and was accompanied by high levels of *FLC* transcript and H3K4me3 modification, as well as low *FT* and *SOC1* transcript levels. However, inducing *FRI* expression at a later stage (i.e., more than one week after germination) did not effectively delay flowering. These data indicate that the timing of *FRI* expression affects flowering time; particularly, the post-embryo and young seedlings are most sensitive to *FRI*-mediated suppression of flowering. Consistent with our study, Sheldon *et al*. found that *FLC* activity during late embryonic development is a prerequisite for the repressive action of *FLC* on flowering (Sheldon *et al*., 2008). Furthermore, *FRI* maintains high levels of *FLC* expression in later embryonic and vegetative development (Choi *et al*., 2009). Reprogramming of *FLC* expression by *FRI* during embryonic and vegetative growth ensures that FT is repressed before winter so that the long-day photoperiod of spring is able to induce *FT* activity, with flowering occurring at an optimal time.

In conclusion, here we reported the spatial and temporal effects of *FRI* on flowering time in *Arabidopsis*, and found spatially expressing *FRI* in the root tissue, not only in the leaves or phloem, also efficiently delayed flowering. We further identified *MAF4* and *MAF5* in the roots acted as the novel target for FRI to delay flowering. Meanwhile, temporally expressing *FRI* during the pro-embryo or post-embryo stage also efficiently activated *FLC* and delay flowering. On the basis of our data, we propose a working model for this mechanism (Fig. 9). Before vernalization, ectopic expression of *FRI* in the embryo, young seedlings, in the leaves or phloem activates *FLC*, or in the roots activates *MAF4* and *MAF5*. Activated *FLC* directly blocks the *FT* signal in the leaves to delay flowering. Correspondingly, activated *MAF4*/*MAF5* in roots might induce the transcription of the gene encoding the antiflorigen-like molecule to antagonize *FT* expression in the leaves. During vernalization, the cold signal could be perceived by the whole plant, relieving the *FT* signal, and finally promoting floral initiation. Taken together, our findings provide insight into the mechanism by which *FRI* modulates flowering during plant development, and demonstrate that roots, and not just leaves, perceive the flowering signal to fine-tune the flowering time.

**Fig. 9.**
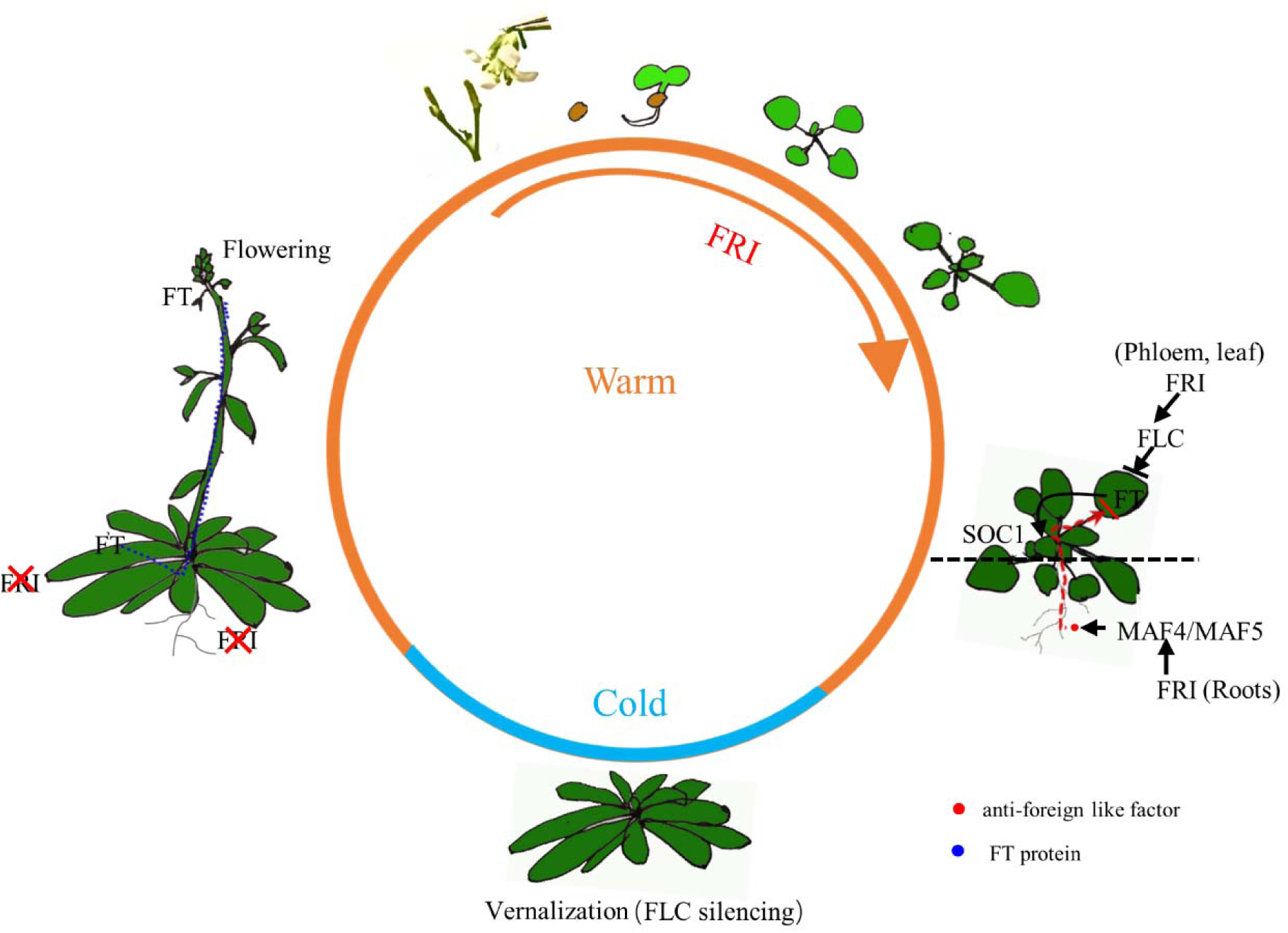
Proposed model in which the spatial-temporal expression of *FRI* modulates flowering time in *Arabidopsis thaliana*. Before vernalization, *FRI* expression in the phloem or leaves could efficiently increase the *FLC* transcripts to suppress the in-situ *FT* signal and delay flowering. Expression of *FRI* in the root could activate *MAF4* and *MAF5*, and possibly generate a certain mobile silencing signal, like anti-florigen-like factor (marked with a red circle), that impairs the *FT* signal in the leaves and finally delays flowering. During vernalization, the cold signal could be perceived by the leaves and root, triggering degradation of *FRI*, which interferes with *FT* activity (marked with a blue circle) and subsequently promotes flowering.

## Supplementary data

**Table S1**. Primers used in this study.

**Fig. S1**. Constitutive expression of *FRI* delays flowering.

**Fig. S2**. Localization of *GUS*-*FRI* expression by GUS staining.

**Fig. S3**. *FRI* expression under the control of the *SUC2*, *RolC*, *KNAT1*, *ML1* and *TobRB7* promoters delayed flowering.

**Fig. S4**. Flowering phenotypes among various transgenic lines after vernalization.

**Fig. S5**. Quantitative RT-PCR analysis of the spatial expressions of *FLC*, *FT* and *SOC1* in different tissues from different transgenic lines after vernalization.

**Fig. S6**. CHIP assay of the relative levels of H3K4me3 in *FLC* chromatin after vernalization.

**Fig. S7**. *MAF1*, *MAF2* or *MAF3* expression in the leaves and root tissue of these transgenic lines was no obvious change.

**Fig. S8**. *MAF4* and *MAF5* expression under the control of the *TobRB7* promoter delayed flowering.

**Fig. S9**. The different flowering times of *ft*, *flc* mutants and the different grafts with *TobRB7::GUS*-*FRI* as the rootstock.

**Fig. S10**. Flowering phenotypes of *ProLEC2::GUS*-*FRI* and *ProEM1::GUS*-*FRI* transgenic lines.

**Fig. S11**. Identification of *ProER8:GUS*-*FRI* by GUS staining.

**Fig. S12**. Quantitative RT-PCR analysis of the expressions of *FT* and *SOC1* in the *ProER8:GUS*-*FRI* line treated with or without estradiol.

## Acknowledgements

We thank Dr. Hailong An for providing the constructs containing different promoters, and ABRC for providing the T-DNA insertion seeds. This article was supported by the National Science Foundation of China (No. 31470348), the Natural Science Foundation of Yunnan Province (NO. 2016FA015) and start-up funding from Shanghai University. X.K. was supported by the National Science Foundation of China (No. 31500221) and the 13th Five-year Informatization Plan of Chinese Academy of Sciences (No. XXH13506).

---

**Supplemental Table 1.**
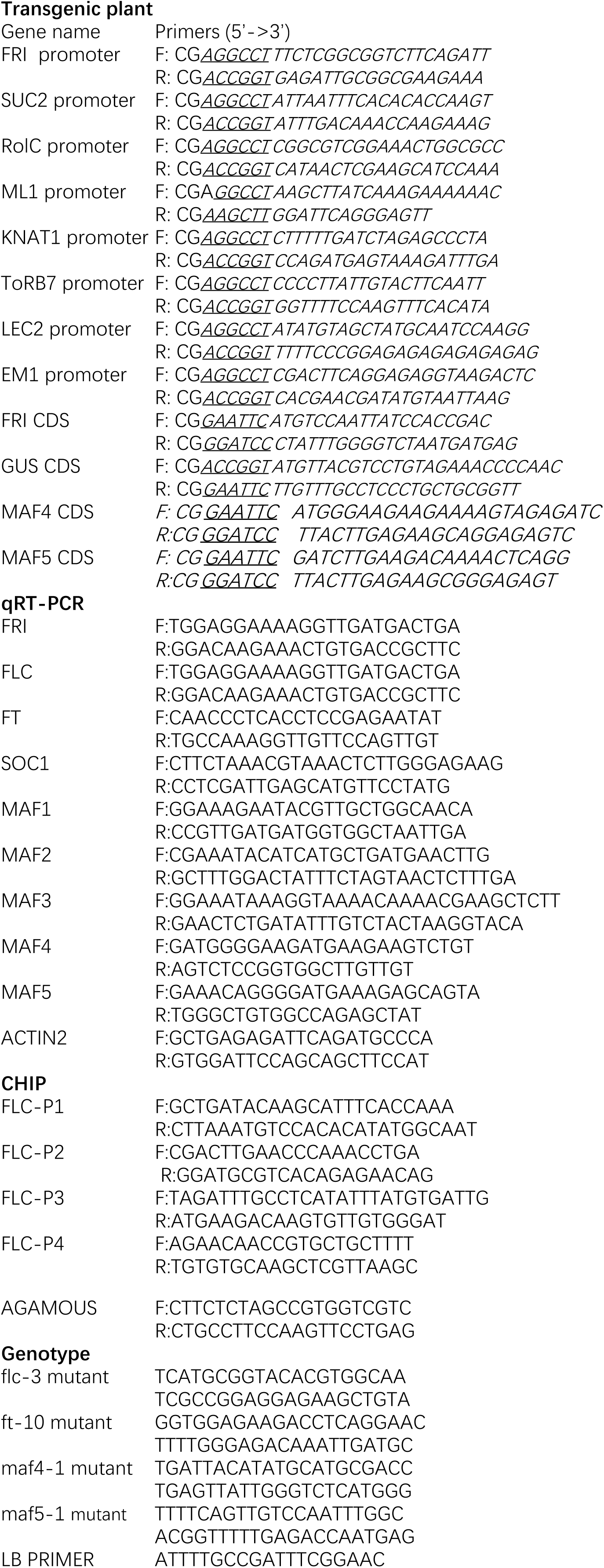
Primers used in this study. Note that sequences underlined indicate the restriction enzyme recognize sequences

